# MB-SupCon: Microbiome-based predictive models via Supervised Contrastive Learning

**DOI:** 10.1101/2022.06.23.497232

**Authors:** Sen Yang, Shidan Wang, Yiqing Wang, Ruichen Rong, Jiwoong Kim, Bo Li, Andrew Y. Koh, Guanghua Xiao, Qiwei Li, Dajiang Liu, Xiaowei Zhan

## Abstract

Human microbiome consists of trillions of microorganisms. Microbiota can modulate the host physiology through molecule and metabolite interactions. Integrating microbiome and metabolomics data have the potential to predict different diseases more accurately. Yet, most datasets only measure microbiome data but without paired metabolome data. Here, we propose a novel integrative modeling framework, Microbiome-based Supervised Contrastive Learning Framework (MB-SupCon). MB-SupCon integrates microbiome and metabolome data to generate microbiome embeddings, which can be used to improve the prediction accuracy in datasets that only measure microbiome data. As a proof of concept, we applied MB-SupCon on 720 samples with paired 16S microbiome data and metabolomics data from patients with type 2 diabetes. MB-SupCon outperformed existing prediction methods and achieves high average prediction accuracies for insulin resistance status (84.62%), sex (78.98%), and race (80.04%). Moreover, the microbiome embeddings form separable clusters for different covariate groups in the lower-dimensional space, which enhances data visualization. We also applied MB-SupCon on a large inflammatory bowel disease study and observed similar advantages. Thus, MB-SupCon could be broadly applicable to improve microbiome prediction models in multi-omics disease studies.

## 1 Introduction

The human microbiome is a collection of living microorganisms cohabitating in distinct body niches [1, 2]. The microbiome significantly impacts human health, including diseases and treatments [3]. Accordingly, it is possible to use microbiome measurements to predict host physiologic conditions non-invasively. Creating microbiome-based prediction models has great benefits for medical research [4].

Earlier work on microbiome-based prediction models using microbiome abundances includes random forest, support vector machines models[5]. While identification and quantification of microbiome taxa using microbiome data alone lead to associative and correlative insights, multi-omics can offer mechanistic insights and potentially improve prediction accuracy over models based on microbiomes alone. For example, in colorectal cancer, specific bacterial species has been associated with increased disease risk [6]. Follow-up mechanistic studies further elucidated the functions of the pathogenic species through multi-omics data analysis [7, 8]. Similar multi-omics approaches, especially in microbiome and metabolomics, have been applied to other diseases [9, 10]. To leverage multi-omics data features and unleash the potential of non-invasive microbiome biomarkers, we aim to develop a general framework for phenotype prediction using microbiome data.

Statistical learning and artificial intelligence research have advanced microbiome-based prediction models. Earlier work utilized taxonomic abundance data and linear or logistic regression models with penalties (e.g., LASSO model, and elastic net model) [11]. More recent approaches integrate multi-omics data using partial least squares (PLS), partial least squares-discriminant analysis (PLS-DA), or canonical correlation analysis (CCA) [12]. These models rely on linear transformations of original features in supervised or unsupervised learning. Recently, contrastive learning has been introduced in the analysis of the multi-omics data [13] that can capture non-linear relationships between features. For example, a simple framework for unsupervised contrastive learning (simCLR) achieves state-of-the-art prediction performance [14]. Supervised contrastive learning (SupCon) in computer vision tasks also demonstrated superior robustness and prediction accuracies [15], and these advantages have solid theoretical support [16]. Inspired by the success of these approaches, we propose a novel supervised-learning framework (MB-SupCon) based on non-linear transformations of multi-omics datasets, which achieve robust and accurate prediction performance. Our method architecture is intuitive and requires only modest-sized multi-omics data. We demonstrate MB-SupCon’s utility using data from a published type 2 diabetes study where MB-SupCon-based model improves prediction accuracies by a large margin; Another independent application of MB-SupCon to an Inflammatory Bowel Disease (IBD) study also produced consistent improvements. Moreover, we demonstrated that the microbiome embeddings from MB-SupCon can better separate different phenotype groups and lead to more informative visualizations of the data. We posit that our microbiome-based prediction model can easily be applied to other disease types and used to integrate data from a variety of omics technologies.

## 2 Results

### 2.1 MB-SupCon: Microbiome-based prediction model via supervised contrastive learning

The main goal of MB-SupCon is to improve the prediction of phenotype or clinical covariates via supervised contrastive learning. An overall workflow is shown in **Figure 1**. The model input includes gut microbiome and metabolome data, phenotype information and/or clinical covariates. We then train a supervised contrastive learning model to obtain the weights of the encoder networks. Finally, we apply the predictive model to independent test datasets to assess its accuracy. The microbiome embedding is critically useful for downstream analysis tasks, including 1) predicting phenotypic outcomes and covariates and 2) visualizing the lower-dimensional representation. We show that approaches using microbiome embedding from MB-SupCon often have better performance than approaches using raw microbiome abundances.

**Figure 1.**
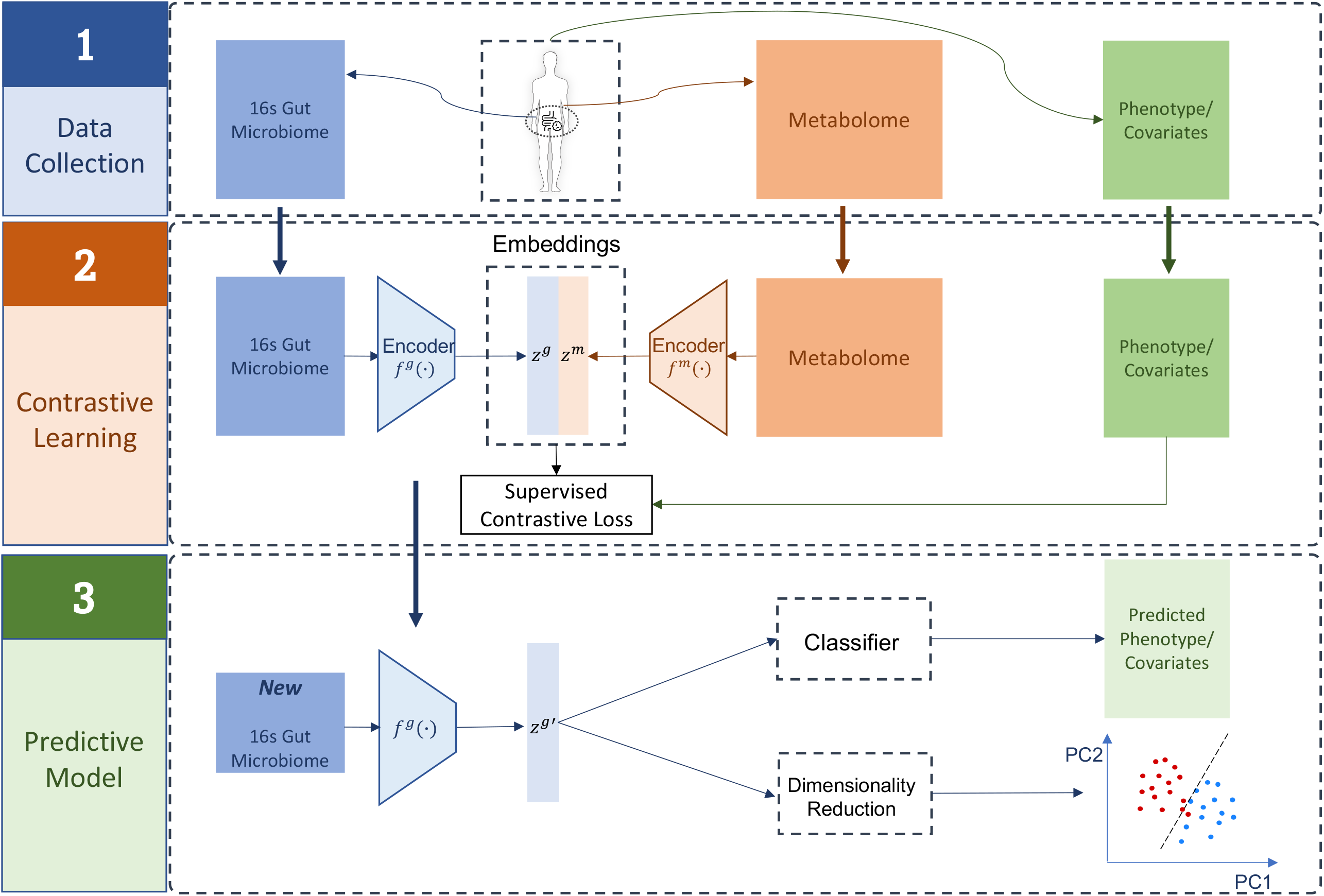
Overview of the MB-SupCon framework. Step 1 - Data Collection: Microbiome, metabolome, phenotype/covariates are collected; Step 2 – Contrastive Learning – MB-SupCon is applied, and two encoder networks are trained; Step 3 – Predictive Model – microbiome encoder network can be applied to new microbiome data to obtain microbiome embeddings. The embeddings lead to an improved microbiome-based prediction model and lower-dimensional representation.

### 2.2 MB-SupCon improved categorical outcome prediction in type 2 diabetes study

We trained MB-SupCon using real human gut microbiome and metabolome data obtained in a host-microbe dynamics study by Zhou, Sailani [17]. The omics data were collected longitudinally from subjects with prediabetes over approximately four years. Gut microbiome data were obtained from stool samples, and host metabolome data was obtained from blood samples at each visit of subjects. We subset both datasets and retained 720 samples with both 16s gut microbiome and metabolome data. Microbiome data is encoded as a matrix of 720 × 96 dimension with entries having values between [0,1), (i.e., [0,1)^720×96^), and each of the 96 features represents the relative abundance of one microbial taxon from 5 taxonomic levels - phylum, class, order, family, and genus. Metabolome data is encoded as a matrix of dimension 720 × 724, with each entry taking values from non-negative real numbers, (i.e.,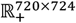), and each of the 724 features represents the abundance of one metabolite. Standardization was applied to both datasets before model fitting so that each feature has a mean value of zero and unit variance. In addition, at each visit, demographic or clinical covariates (e.g., sex, age, insulin resistant/insulin sensitive, BMI, etc.) were also recorded for all subjects. We also attempted to predict the covariates using microbiome and metabolome data to evaluate different predictive models. To evaluate the predictive performance for each machine learning model, we applied 12 random splitting of training (70%), validation (15%), and testing (15%) to the data. For each split, the training and validation sets were used for model fitting and hyperparameter tuning (**Supplementary texts: Training and tuning procedure**), and the testing set was used for benchmarking.

To illustrate the advantage of MB-SupCon, we used

- a logistic regression with elastic net regularization (EN),
- a multi-layer perceptron (MLP),
- a support vector machine classifier (SVM),
- a random forest classifier (RF)

to analyze and compare their performance on

- the original microbiome abundances,
- the embedding of supervised contrastive learning (MB-SupCon).

We also compared MB-SupCon with a method that uses a logistic regression model to analyze unsupervised embeddings (MB-simCLR).

To distinguish analyses using original abundance and embeddings, we denote methods that analyze embeddings with prefix “MB-SupCon” e.g., MB-SupCon+MLP represents using MLP to analyze MB-SupCon embeddings.

We listed the details in **Supplementary texts: Calculation of the microbiome embedding** on obtaining microbiome embeddings in unsupervised or supervised learning. To evaluate prediction accuracy, we compute the fraction of correctly predicted labels for each model. Since we create multiple splits of the data for training, validation, and testing, the average prediction accuracy using different test folds are reported.

MB-SupCon embeddings, compared with the original data, lead to improved prediction accuracies in logistic regression with an elastic net penalty, SVM, MB-simCLR. The methods using MB-SupCon embedding almost always outperform RF and MLP models using raw microbiome abundance, which are two of the most accurate methods (**Table 1, Figure 3**). For the prediction of insulin resistance, methods using MB-SupCon embeddings achieved 84.62% average accuracy (MB-SupCon+Logistic, MB-SupCon+SVM, MB-SupCon+RF, and MB-SupCon+MLP), which is better than methods that uses raw abundances, i.e., the elastic net logistic regression (76.69%), SVM (79.46%), MB-simCLR (65.67%), and similar to RF (83.93%) and MLP (83.73%). Similarly, for predicting sex, MB-SupCon also has good average prediction accuracy (78.98%). For predicting race, a four-category outcome, approaches using MB-SupCon embeddings reaches the lead average accuracy (80.04%), and their advantage is consistent over the other methods, including RF (77.90%) and MLP (75.60%). More importantly, MB-SupCon embedding leads to a near-best prediction accuracy regardless of the choice of machine learning algorithms, which demonstrated its utility and robustness.

**Table 1.** Average prediction accuracies on testing data from 12 random training-validation-testing splits, by using different methods for categorical covariates (T2D study). Acronyms: Logistic - logistic regression with elastic net penalty using original data; SVM - support vector machine classifier using original data; RF - random forest classifier using original data; MLP multi-layer perceptron using original data; MB-simCLR - logistic regression model with elastic net penalty using microbiome embeddings learned from unsupervised contrastive learning; MB-SupCon + Logistic - logistic regression model with elastic net penalty using microbiome embeddings learned from supervised contrastive learning. MB-SupCon + SVM: support vector machine classifier using microbiome embeddings learned from supervised contrastive learning; MB-SupCon + RF: random forest classifier using microbiome embeddings learned from supervised contrastive learning; MB-SupCon + MLP: multi-layer perceptron using microbiome embeddings learned from supervised contrastive learning; Avg. Acc. based on MB-SupCon: average accuracies among MB-SupCon + Logistic, MB-SupCon + SVM, MB-SupCon + RF and MB-SupCon + MLP.

| **Prediction Task** | **Logistic** | **SVM** | **RF** | **MLP** | **MB-simCLR** |
| --- | --- | --- | --- | --- | --- |
| Insulin resistance | 76.69% | 79.46% | 83.93% | 83.73% | 65.67% |
| Sex | 65.61% | 69.02% | 80.38% | 78.94% | 59.85% |
| Race | 72.99% | 72.17% | 77.90% | 75.60% | 68.38% |

| **Prediction Task** | **MB-SupCon + Logistic** | **MB-SupCon + SVM** | **MB-SupCon + RF** | **MB-SupCon + MLP** | **Avg. Acc. based on MB-SupCon** |
| --- | --- | --- | --- | --- | --- |
| Insulin resistance | 84.42% | 85.12% | 84.62% | 84.33% | 84.62% |
| Sex | 78.94% | 79.02% | 79.24% | 78.71% | 78.98% |
| Race | 80.73% | 80.36% | 79.91% | 79.17% | 80.04% |

### 2.3 MB-SupCon better visualized embeddings in independent datasets

In addition to improving prediction accuracy, MB-SupCon embeddings in the lower dimensional space can be useful for visualizations. In **Figure 2A**, we applied PCA on 1) raw abundance data, 2) embeddings from MB-simCLR, and 3) embeddings from MB-SupCon in an independent test data. We placed the samples of test datasets onto the principal component 2 (PC2) vs 1 (PC1) scatterplot using a random seed of 1. In addition, we also compared MB-SupCon to three other methods, i.e., Sparse Partial Least Squares Discriminant Analysis (sPLS-DA) [18], Sparse Partial Least Squares (sPLS)[19], and Data Integration Analysis for Biomarker discovery using Latent cOmponents (DIABLO) [20], for their capability to distinguish different groups of covariates. sPLS-DA [18] predicts covariates using microbiome data only; the other two methods are based on integrative modeling of both microbiome and metabolome data. sPLS [19] uses microbiome data as predictors and metabolome data as responses. DIABLO [20] uses multiple omics data from the same samples to be blocks and covariate values to be the outcome. All three methods can be implemented under the “mixOmics” [12] framework. In **Figure 2B**, we compared the lower-dimensional scatterplots (Component 2 vs. Component 1) on the same testing data for each method to those of MB-SupCon in **Figure 2A**. Only the embedding from MB-SupCon leads to separable clusters from distinct covariate groups, whereas the other established methods failed to separate different categories of covariates. This result confirms that the improvements in prediction accuracy of MB-SupCon can be attributable to better feature embeddings.

**Figure 2.**
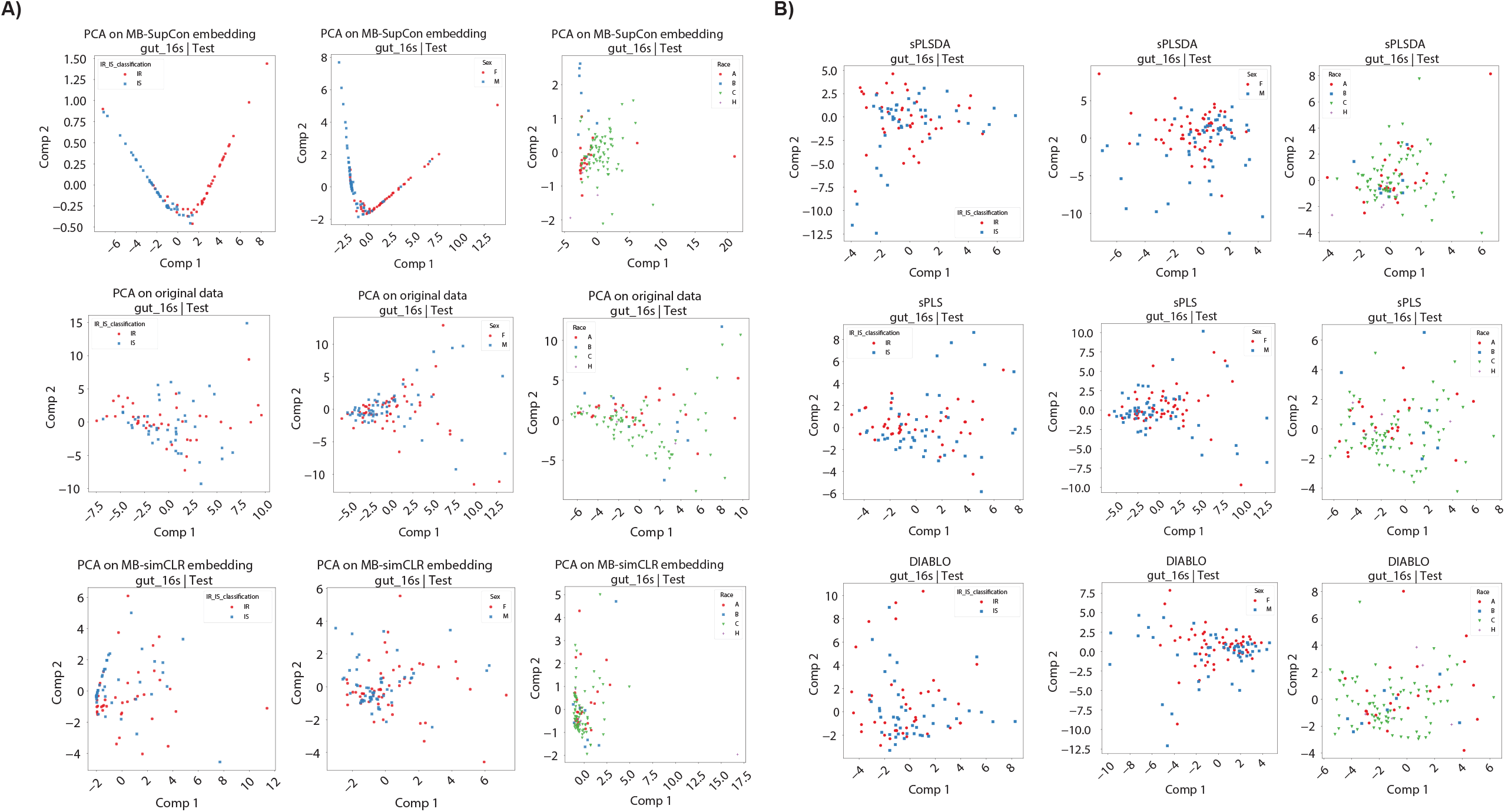
Scatter plots of test data on lower-dimensional space (T2D study). **Panel A: Scatter plots of test data (random seed 1) on a 2-dimensional space by PCA.** 1st row: the first two principal components for the embeddings learned from MB-SupCon; 2nd row: the first two principal components for the original data; 3rd row: the first two principal components for the embeddings learned from MB-simCLR. Acronyms: PCA - Principal component analysis. **Panel B: Scatter plots of test data (random seed 1) on 2-dimensional space by other methods**. 1st row: the first two components learned from sPLSDA on original data; 2nd row: the first two components learned from sPLS on original data; 3rd row: the first two principal components learned from DIABLO on original data. Acronyms: PCA - Principal component analysis. sPLS-DA - Sparse Partial Least Squares Discriminant Analysis; sPLS - Sparse Partial Least Squares; DIABLO - Data Integration Analysis for Biomarker discovery using Latent cOmponents.

**Figure 3.**
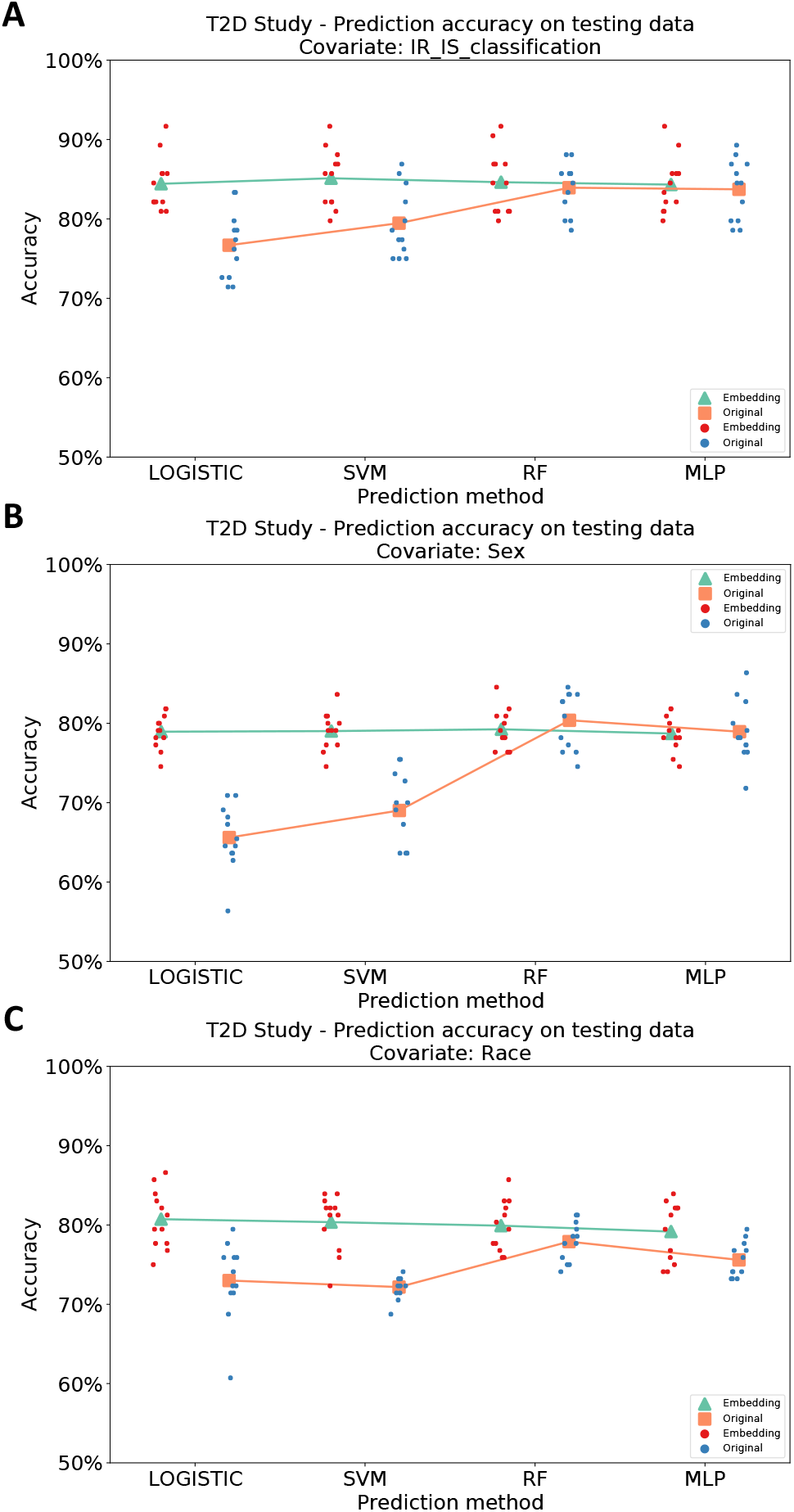
Scatter plot of average prediction accuracies on test data from 12 random training-validation-testing splits, by using different methods for categorical covariates (T2D study). Green triangles and red points represent predictions based on MB-SupCon embeddings. Orange squares and blue points represent predictions based on original microbiome data. **Panel A**: Insulin resistant/sensitive; **Panel B**: Sex**; Panel C**: Race. Acronyms: LOGISTIC **-** logistic regression with elastic net penalty; SVM - support vector machine classifier; RF - random forest classifier; MLP - multi-layer perceptron.

### 2.4 MB-SupCon analysis of an inflammatory bowel disease study

To further evaluate the performance of MB-SupCon, we applied it to another independent multi-omics Inflammatory Bowel Disease (IBD) study with both metagenomics and metabolomics data [9] (detailed in **Supplementary texts: Network architecture and training of MB-SupCon model for IBD study)**. With “diagnosis” of IBD status as the covariate, we trained, validated and tested our model using 12 different random splits similar to the diabetes study. For each model, we evaluated the model performance on testing data. As shown in **Table 2** and **Figure 4**, the results remained consistent with the T2D study. Approaches using MB-SupCon embeddings achieved significantly better average prediction accuracies (74.04%) compared to approaches using original data directly, including logistic regression (67.79%) and SVM (52.70%). When RF or MLP is used, predictions based on MB-SupCon embedding was comparable to the predictions using original abundance information, although MB-SupCon+RF had a slightly smaller variance compared to RF and has a marginal advantage compared to MLP. This validated the reliability and extensive applicability of MB-SupCon.

**Table 2.**
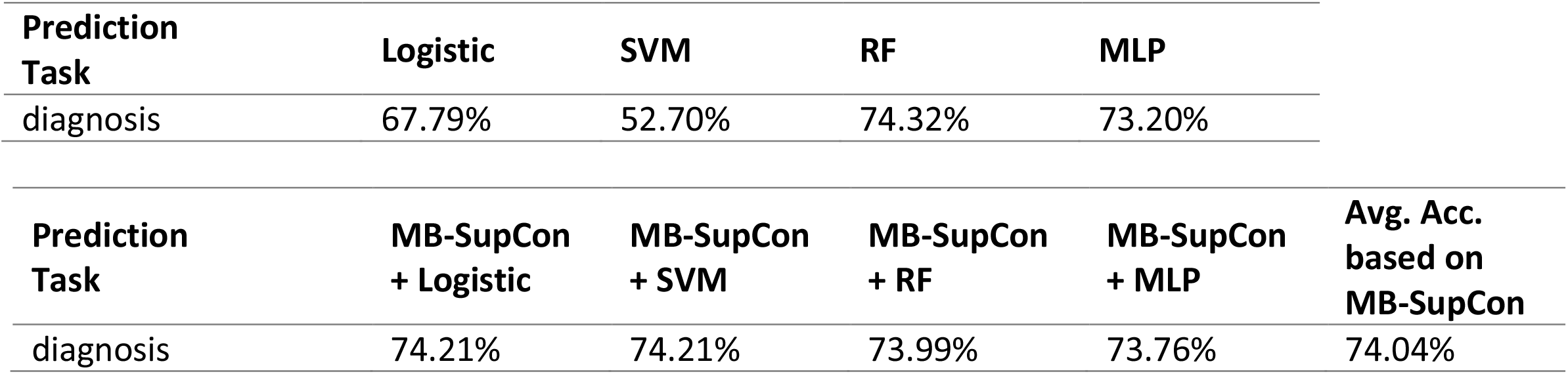
Average prediction accuracies on testing data from 12 random training-validation-testing splits, by using different methods for categorical covariates (IBD study). Acronyms are defined the same as those from Table 1.

| **Prediction Task** | **Logistic** | **SVM** | **RF** | **MLP** |
| --- | --- | --- | --- | --- |
| diagnosis | 67.79% | 52.70% | 74.32% | 73.20% |

| **Prediction Task** | **MB-SupCon + Logistic** | **MB-SupCon + SVM** | **MB-SupCon + RF** | **MB-SupCon + MLP** | **Avg. Acc. based on MB-SupCon** |
| --- | --- | --- | --- | --- | --- |
| diagnosis | 74.21% | 74.21% | 73.99% | 73.76% | 74.04% |

**Figure 4.**
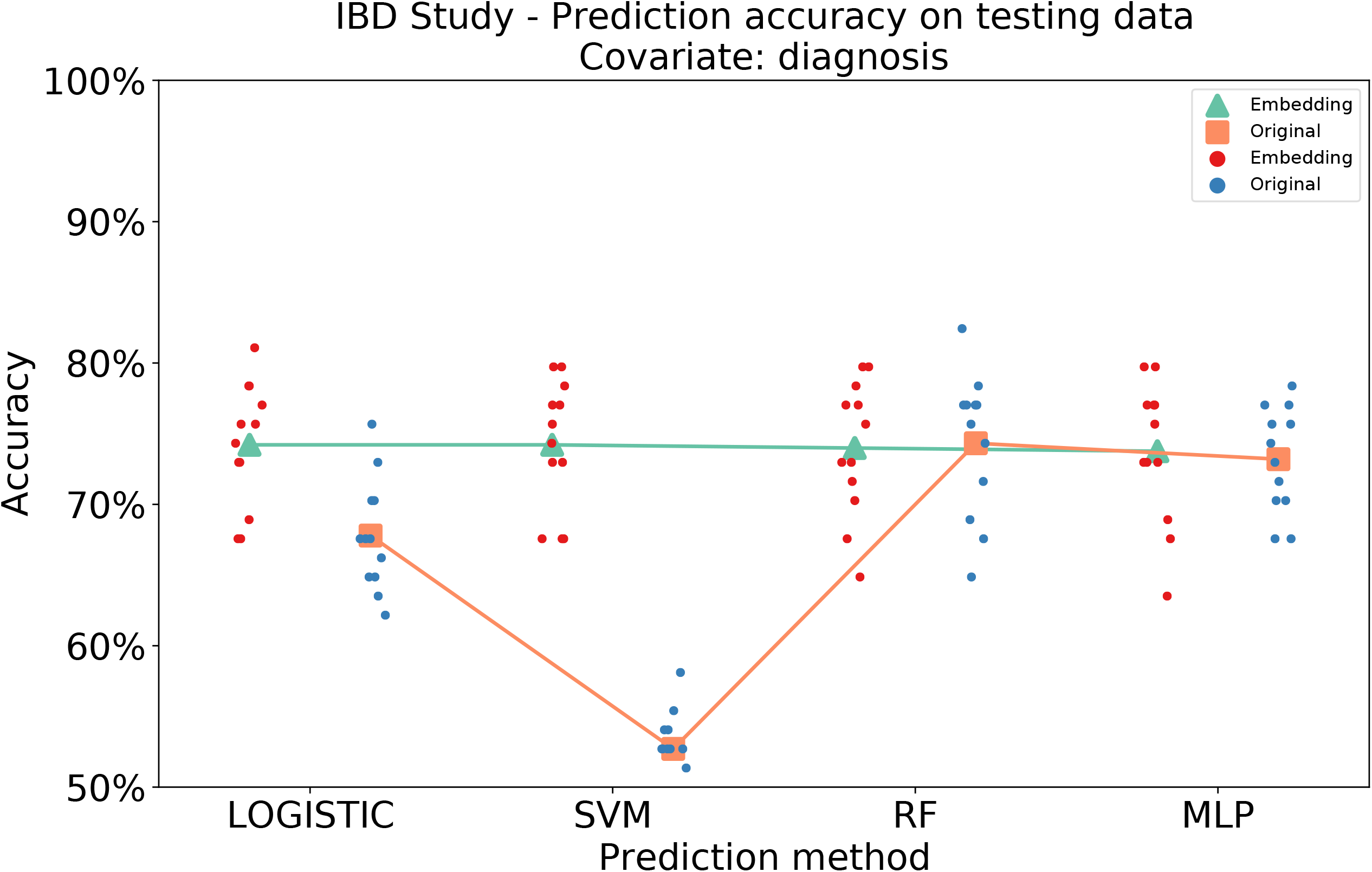
Scatter plots of average prediction accuracies for diagnosis on testing data from 12 random training-validation-testing splits, by using different methods for categorical covariates (IBD study). Green triangles and red points represent predictions based on MB-SupCon embeddings. Orange squares and blue points represent predictions based on original microbiome data. Acronyms: LOGISTIC **-** logistic regression with elastic net penalty; SVM - support vector machine classifier; RF - random forest classifier; MLP - multi-layer perceptron.

## 3 Discussion

A reliable microbiome-based prediction model could have immediate values in disease diagnosis and treatment responses prediction [6, 21, 22]. Here, we propose a novel method, MB-SupCon, to improve those models by utilizing increasingly accessible multi-omics datasets. The method leverages the strengths of contrastive learning, which were first established in computer vision tasks [14-16, 23]. MB-SupCon performs the nonlinear transformation of microbiome abundance and produces useful embeddings, which lead to improved prediction accuracies and more informative visual representations. We demonstrate these advantages of MB-SupCon utilizing existing published data from a diabetes study and an inflammatory bowel disease study. We showed that the improved microbiome prediction model using MB-SupCon embeddings is more accurate than elastic net logistic regression, support vector machine, and unsupervised contrastive learning model, and can achieve comparable or better performance of random forest and multi-layer perceptron.

Like all other deep learning models, MB-SupCon has limitations. One drawback is that it does not explicitly offer biological interpretations between the microbiome and metabolomics. This “black-box” nature of the deep learning model often leads to criticisms. Developing more interpretable machine learning models can potentially address the emerging biological questions. Another limitation is that MB-SupCon does not explicitly model sample relatedness. Specifically, as paired longitudinal data is relatively infrequent, MB-SupCon does not incorporate features that could account for correlations among longitudinal samples. A better solution to explore in the future is to change the current MLP encoders to mixed effect neural networks [24, 25] so that variation within subjects for longitudinal data could be better modeled and explained.

There are numerous future applications and extensions of MB-SupCon. MB-SupCon is not restricted to the microbiome and metabolomic data analysis. It can be applied to any omics technology (e.g., proteomics, host transcriptomics, host methylome, etc.). Moreover, MB-SupCon can be extended to integrate more than two types of omics data. This can be achieved by adding pair-wise supervised contrastive losses.

In summary, we believe MB-SupCon and encoder-based on the neural network in general have advantage in approximating non-linear functions and modeling high-dimensional data. MB-SupCon framework can be applicable in broad multi-omics settings and improves microbiome-based prediction models.

## 4 Methods

Contrastive learning aims to maximize the similarities between microbiome embedding and metabolome embedding from a pair of samples. Let *X*^*g*^ and *X*^*m*^ be the standardized microbiome and metabolome data. Suppose there are *n* samples in a minibatch. For a single sample *i* (*i* = 1, 2, …, *n*), we denote the associated microbial and metabolic data as 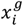 and 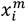, respectively. Let the microbiome (or metabolome) encoder network be a multi-layer perceptron *f*^*g*^(·) (or *f*^*m*^(·)). The encoded features (embeddings) of microbiome and metabolome for sample *i* are 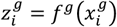 and 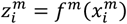, respectively. We define the similarity between the encoded vectors 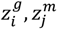 for *i, j*∈ S = {1,2, …, *n*} in the latent space by the cosine similarity,

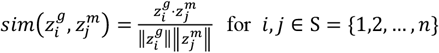

where · denotes the dot product of two vectors and || || denotes the Euclidean norm of a vector.

We first introduce MB-SimCLR, an unsupervised contrastive learning approach: if a pair of microbiome and metabolome samples are from the same sample, we define the corresponding data 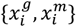 as a “positive pair”. Otherwise, we define the pair of data 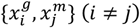 as a “negative pair”. Given *n* pairs of microbiome and metabolome samples, if we set the embedding vector of microbiome 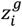 as an anchor, the loss of unsupervised contrastive learning is

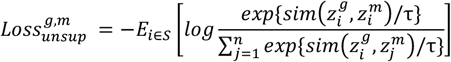

where *i* ∈ *S* = {1,2, …, *n*}, τ ∈ ℝ _+_ is the temperature parameter.

Symmetrically, by anchoring the embedding of the metabolome we can get loss 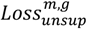. The total loss will be the sum of these two parts: 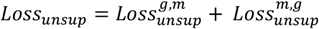

Improved upon MB-SimCLR, we describe a supervised contrastive learning method, MB-SupCon, where we incorporate labels in calculating the loss function. Given a specific categorical label *y*_*i*_ from sample *i, P*(*y*_*i*_) denotes the index set of samples with label *y*_*i*_. Any pairs of microbiome and metabolome vectors 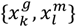 with *K, l*∈ *P*(*y*_*i*_) are treated as “positive pairs”. Otherwise, they are “negative pairs”. Suppose we set microbiome embedding 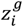 for *i* ∈ *S* with label *y*_*i*_ as an anchor. Then supervised contrastive loss [15] is defined as

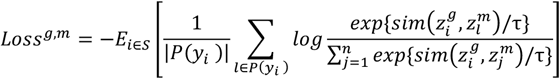

where |*P*(*y*_*i*_)| is the cardinality of index set *P*(*y*_*i*_), τ ∈ ℝ_+_ is the temperature parameter.

By anchoring metabolome embedding, we can get *Loss*^*m,g*^. The total loss is still the sum of *Loss*^*m,g*^ and *Loss*^*m,g*^.

In all, the difference between supervised contrastive learning and unsupervised contrastive learning is the definition of positive and negative sample pairs. Once the loss is determined, we can update the weights of encoder networks using the stochastic gradient descent (SGD) method. Embedding can be calculated as the network outputs. Details are provided in the **Supplemental Texts: Network Architecture and Training**.

## Figure and table legend

**Supplementary Figure 1. Structure of the microbiome and metabolome encoder network**. Only dense layers are visualized where numbers represent neuron counts. Batch normalized layer, Activation layer, Dropout layer are appended after each dense layer but not shown.

**Supplementary Figure 2. Hyperparameter tuning for MB-SupCon on T2D study**. Panel A1 – A3: hyperparameter tuning result for covariate Insulin resistant/sensitive by logistic regression with an elastic net penalty, SVM, and RF; Panel B1 - B3: hyperparameter tuning results for covariate Sex by logistic regression with an elastic net penalty, SVM, and RF; Panel C1 – C3: hyperparameter tuning result for covariate Race by logistic regression with an elastic net penalty, SVM and RF.

Acronyms: Dropout Rate: dropout rate of the encoders from MB-SupCon; weight_decay: weight decay value (l2 regularization) of the stochastic gradient descent (SGD) optimizer; temperature: temperature hyperparameter when calculating contrastive losses.

## Reference

[1] Human Microbiome Project C. A framework for human microbiome research. Nature. 2012;486:215–21.

[2] Sender R, Fuchs S, Milo R. Revised Estimates for the Number of Human and Bacteria Cells in the Body. PLoS Biol. 2016;14:e1002533.

[3] Cho I, Blaser MJ. The human microbiome: at the interface of health and disease. Nature reviews Genetics. 2012;13:260–70.

[4] Claesson MJ, Clooney AG, O’Toole PW. A clinician’s guide to microbiome analysis. Nat Rev Gastroenterol Hepatol. 2017;14:585–95.

[5] Pasolli E, Truong DT, Malik F, Waldron L, Segata N. Machine Learning Meta-analysis of Large Metagenomic Datasets: Tools and Biological Insights. PLoS Comput Biol. 2016;12:e1004977.

[6] Yu J, Feng Q, Wong SH, Zhang D, Liang QY, Qin Y, et al. Metagenomic analysis of faecal microbiome as a tool towards targeted non-invasive biomarkers for colorectal cancer. Gut. 2017;66:70–8.

[7] Chen F, Dai X, Zhou CC, Li KX, Zhang YJ, Lou XY, et al. Integrated analysis of the faecal metagenome and serum metabolome reveals the role of gut microbiome-associated metabolites in the detection of colorectal cancer and adenoma. Gut. 2021.

[8] Wang Q, Ye J, Fang D, Lv L, Wu W, Shi D, et al. Multi-omic profiling reveals associations between the gut mucosal microbiome, the metabolome, and host DNA methylation associated gene expression in patients with colorectal cancer. BMC Microbiol. 2020;20:83.

[9] Lloyd-Price J, Arze C, Ananthakrishnan AN, Schirmer M, Avila-Pacheco J, Poon TW, et al. Multi-omics of the gut microbial ecosystem in inflammatory bowel diseases. Nature. 2019;569:655–62.

[10] Heintz-Buschart A, May P, Laczny CC, Lebrun LA, Bellora C, Krishna A, et al. Integrated multi-omics of the human gut microbiome in a case study of familial type 1 diabetes. Nat Microbiol. 2016;2:16180.

[11] Friedman J, Hastie T, Tibshirani R. The elements of statistical learning: Springer series in statistics Springer, Berlin; 2001.

[12] Rohart F, Gautier B, Singh A, Lê Cao K-A. mixOmics: An R package for ‘omics feature selection and multiple data integration. PLOS Computational Biology. 2017;13:e1005752.

[13] Tian Y, Krishnan D, Isola P. Contrastive multiview coding. European conference on computer vision: Springer; 2020. p. 776–94.

[14] Chen T, Kornblith S, Norouzi M, Hinton G. A simple framework for contrastive learning of visual representations. International conference on machine learning: PMLR; 2020. p. 1597–607.

[15] Khosla P, Teterwak P, Wang C, Sarna A, Tian Y, Isola P, et al. Supervised contrastive learning. Advances in Neural Information Processing Systems. 2020;33:18661--73.

[16] Tian Y, Krishnan D, Isola P. Contrastive Multiview Coding. European conference on computer vision. 2020:776--94.

[17] Zhou W, Sailani MR, Contrepois K, Zhou Y, Ahadi S, Leopold SR, et al. Longitudinal multiomics of host–microbe dynamics in prediabetes. Nature. 2019;569:663–71.

[18] Le Cao KA, Boitard S, Besse P. Sparse PLS discriminant analysis: biologically relevant feature selection and graphical displays for multiclass problems. BMC Bioinformatics. 2011;12:253.

[19] Lê Cao K-A, Rossouw D, Robert-Granié C, Besse P. A sparse PLS for variable selection when integrating omics data. Statistical applications in genetics and molecular biology. 2008;7.

[20] Singh A, Shannon CP, Gautier B, Rohart F, Vacher M, Tebbutt SJ, et al. DIABLO: an integrative approach for identifying key molecular drivers from multi-omics assays. Bioinformatics. 2019;35:3055–62.

[21] Andrews MC, Duong CPM, Gopalakrishnan V, Iebba V, Chen WS, Derosa L, et al. Gut microbiota signatures are associated with toxicity to combined CTLA-4 and PD-1 blockade. Nat Med. 2021;27:1432–41.

[22] Zeller G, Tap J, Voigt AY, Sunagawa S, Kultima JR, Costea PI, et al. Potential of fecal microbiota for early-stage detection of colorectal cancer. Mol Syst Biol. 2014;10:766.

[23] Wu Z, Xiong Y, Yu S, Lin D. Unsupervised Feature Learning via Non-Parametric Instance-level Discrimination. Proceedings of the IEEE conference on computer vision and pattern recognition. 2018:3733--42.

[24] Xiong Y, Kim HJ, Singh V. Mixed effects neural networks (menets) with applications to gaze estimation. Proceedings of the IEEE/CVF Conference on Computer Vision and Pattern Recognition 2019. p. 7743–52.

[25] Tandon R, Adak S, Kaye JA. Neural networks for longitudinal studies in Alzheimer’s disease. Artificial intelligence in medicine. 2006;36:245–55.

