## Supplemental Material for "MB-SupCon: Microbiome-based predictive models via Supervised Contrastive Learning"

### Supplemental Texts

#### 1. Network architecture, training, and tuning of MB-SupCon models for T2D study

##### 1.1 Overview

We implemented MB-SupCon, a supervised contrastive learning model for microbiome data and metabolomics data. In the supplemental texts, we provide additional implementation details on the network architecture and training process. The source codes are also available at: https://github.com/ya61sen/MB-SupCon .

##### 1.2 Encoder networks

We designed encoder networks for microbiome and metabolome data, denoted as microbiome encoder $f^g(\cdot)$ and metabolome encoder $f^m(\cdot)$ (**Figure 1**, **Supplementary Figure 1**). The network architecture is similar to a 3-layer perceptron network. After each fully connected layer, we added batch normalization layers to improve stability and dropout layers to remedy overfitting.

##### 1.3 Training and tuning procedure

We used 12 random seeds to randomly split the paired 16S gut microbiome and metabolome data into training dataset (70%), validation dataset (15%), and test dataset (15%). For each split, we trained three supervised contrastive learning models for three tasks: insulin resistant, sex, and race. For each model, we removed data rows with missing labels. The main differences among these three models were the covariates used for prediction labels, which means different supervised contrastive losses were calculated in training, and different encoder networks were obtained.

We set 1,000 iterations of training with a batch size of 32. The structure of encoders for microbiome and metabolome are described in **Methods** and **Supplementary Figure 1**. We used dropout to reduce overfitting. Stochastic gradient descent (SGD) method with a learning rate of 0.001, a momentum of 0.9, and a weight decay ($l2$ regularization) after tunning was used as the optimizer. To avoid gradient explosion, we also clipped the gradient of the model parameter by restraining the norm of the gradient within $(-3, 3)$ during each iteration.

Model tuning was based on validation datasets. Average accuracy on validation data among all 12 splits was used as the evaluation metric. We tuned the model regarding 3 hyperparameters: dropout rate of the encoder, weight decay of the stochastic gradient descent (SGD) optimizer, and the temperature coefficient in calculating contrastive losses. The value set of each hyperparameter for tuning is as follows.

(1) Dropout rate of the encoder. The set of dropout rate for tuning is {0.2, 0.4, 0.6, 0.8}.
(2) Weight decay (l2 penalty). The set of weight decay for tuning is {0.0025, 0.005, 0.01, 0.02, 0.04, 0.08, 0.16, 0.32}.
(3) Temperature. The set of temperature for tuning is {0.125, 0.25, 0.5, 1, 2}.

For each model, we predicted the covariate based on the embeddings from the validation datasets. We chose one predictor among logistic regression with an elastic net penalty, support vector machine, random forest, and a one-layer MLP. The best combination of hyperparameters was then determined by their average accuracies. Then this best model was used for subsequent prediction tasks. The detailed tuning results are shown in **Supplementary Figure 2.**

For each covariate, the best combinations of hyperparameters are listed as follows.

| Prediction Task | Dropout Rate | Weight decay | Temperature |
|---|---|---|---|
| Insulin resistance | 0.6 | 0.08 | 0.25 |
| Sex | 0.4 | 0.04 | 0.125 |
| Race | 0.2 | 0.01 | 0.125 |

##### 1.4 Key differences to MLP (multi-layer perceptron) model

Here we provide critical differences between the MB-SupCon prediction model and the MLP model. In MB-SupCon, the prediction model utilized the microbiome embedding, and the embedding was then fed to a predictor (among 4 options: logistic regression with an elastic net penalty, support vector machine, random forest, and a one-layer MLP). This may be seen to be very similar to an MLP model at the first glance, especially the encoder part from MB-SupCon. However, the key differences remain. First, the training of the MB-SupCon encoder is governed by supervised contrastive loss, which integrates the information from metabolome data and differs from the usual cross-entropy loss used by MLP. Second, we observe that the MB-SupCon model has better or similar performance in all three prediction models (As shown in **Table 1**, the average accuracies for MLP on original data are 83.73%, 78.94%, and 75.60% for insulin resistance, sex, and race, respectively). Third, MB-SupCon shows smooth loss curves in the training stage. The loss curve decreases as the epoch increases for MB-SupCon. In contrast, the loss curve for MLP often varies at large epochs. Thus, we observe the MB-SupCon model is robust to train probably, which confirms that contrastive learning usually requires fewer training samples.

#### 2. Network architecture and training of MB-SupCon model for IBD study

For the IBD study, we also used MLP as the encoders. For the metagenomics encoder, the architecture is as follows: 1479- 512-128-32-10; for the metabolomics encoder, the architecture is as follows: 5084-2048-512-128-32-10. Here, each number represents neuron counts in each layer. Similarly, we also added batch normalization layers and dropout layers after each fully connected layer.

Twelve random training-validation-testing splits were applied to the paired metagenomics and metabolomics data (70%, 15%, and 15% for training, validation, and testing data, respectively). The model performance was evaluated based on the average prediction accuracy on the testing data from all splits.

We also set 1,000 iterations of training with a batch size of 32. For covariate "diagnosis", the dropout rate was set to 0.4. Stochastic gradient descent (SGD) method with a learning rate of 0.001, a momentum of 0.9, and a weight decay (l2 regularization) of 0.04 was used as the optimizer. We also clipped the gradient of the model parameter by restraining the norm of the gradient within $(-3, 3)$ during each iteration.

#### 3. Calculation of the microbiome embedding

The embeddings (representations) of the microbiome (or metabolome) data can be computed from original microbiome (or metabolome) data, respectively. Suppose the original microbiome data for sample $i$ is $x_i^g$ and the microbiome encoder is $f^g(\cdot)$, then the microbiome embedding is

$z_i^g = f^g(x_i^g)$. It is important to note that the network weights were determined in the training process where no test data were used.

Based on these embedding, we conducted two types of analysis:

(1) build a microbiome-based prediction model for different outcomes. We used one of the predictors among logistic regression with an elastic net penalty, support vector machine, random forest, and a one-layer MLP. All these models were trained on the embedding of the training dataset. We then calculated accuracies from the embedding of the test dataset. The average accuracies on test data from 12 random training-validation-testing splits are reported as the outcome.

(2) visualize the embedding in lower dimensions with colors representing different outcomes (or covariates). We applied principal component analysis (PCA) to the embedding of the test dataset to avoid overfitting. Other dimensionality reduction techniques (e.g., t-SNE, UMAP) were also applicable for data visualization.

#### 4. The implementation of sPLSDA, sPLS, and DIABLO analysis based on the "mixOmics" platform

sPLSDA, sPLS, and DIABLO analysis were all based on the "mixOmics" platform (R package "mixOmics") [1]. To compare the ability in distinguishing clusters of covariate groups, we used the same training-validation-testing split (by random seed 1) as MB-SupCon presented in **Figure 2A**.

sPLSDA [2] predicted covariates using microbiome data only. In R, we used the "splsda" function to build the model. The first two components were included for visualization and all other arguments are kept as default. sPLS [3] used microbiome data as predictors and metabolome data as responses. Covariate information was not used during training. In R, we used the "spls" function to build the model. The first two components were used for visualization and all other arguments are kept as default. DIABLO [4] used multiple omics data from the same samples to be blocks and covariate values to be the outcome. In R, we used the "block.splsda" function to build the model. We assumed microbiome and metabolome data were fully connected and the design matrix was set to be "full". The first two components were used for visualization and all other arguments are kept as default.

### Reference

[1] Rohart F, Gautier B, Singh A, Lê Cao K-A. mixOmics: An R package for ‘omics feature selection and multiple data integration. PLOS Computational Biology. 2017;13:e1005752.
[2] Le Cao KA, Boitard S, Besse P. Sparse PLS discriminant analysis: biologically relevant feature selection and graphical displays for multiclass problems. BMC Bioinformatics. 2011;12:253.
[3] Lê Cao K-A, Rossouw D, Robert-Granié C, Besse P. A sparse PLS for variable selection when integrating omics data. Statistical applications in genetics and molecular biology. 2008;7.
[4] Singh A, Shannon CP, Gautier B, Rohart F, Vacher M, Tebbutt SJ, et al. DIABLO: an integrative approach for identifying key molecular drivers from multi-omics assays. Bioinformatics. 2019;35:3055-62.
