## Supplementary figures and images for "MB-SupCon: Microbiome-based predictive models via Supervised Contrastive Learning"

### Supplementary Figure 1

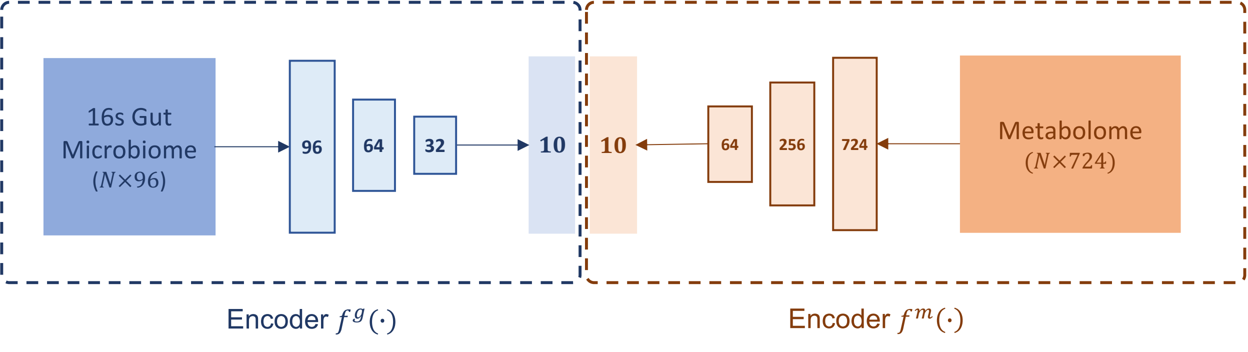

### Supplementary Figure 2

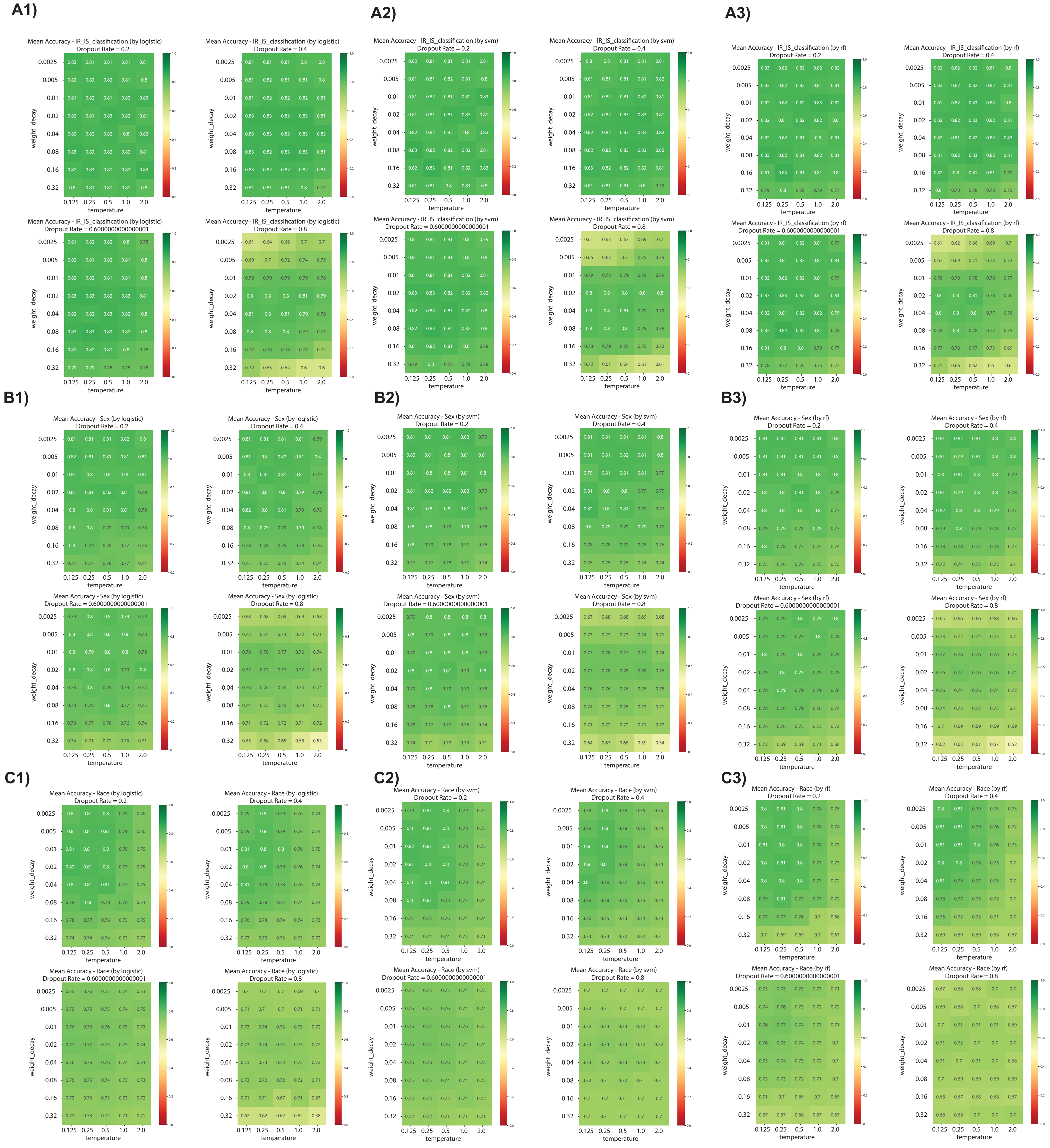
